# Actin cytoskeletal remodeling requires the interaction between Solo and LARG in response to substrate stiffness

**DOI:** 10.1101/2025.04.08.647765

**Authors:** Aoi Kunitomi, Yusuke Toyofuku, Shuhei Chiba, Nahoko Higashitani, Atsushi Higashitani, Shinichi Sato, Kensaku Mizuno, Kazumasa Ohashi

## Abstract

In response to external mechanical stimuli, cells remodel their actin cytoskeleton. Solo, a Rho guanine nucleotide exchange factor (RhoGEF), is involved in mechanical stress responses. Using BioID, we identified PDZ-RhoGEF (PRG), a member of the RGS-RhoGEF family (regulator of G protein signaling domain-containing RhoGEFs, as a Solo-interacting protein. Moreover, we found that Solo regulates PRG during the mechanical stress response. Furthermore, we identified leukemia-associated RhoGEF (LARG), another RGS-RhoGEF member, as a Solo-interacting protein; however, the functional role of this interaction remains unknown. Therefore, in this study, we investigated the interaction between Solo and LARG and found that LARG localizes to Solo accumulation sites at the basal plane and that LARG is required for Solo-induced actin polymerization. Additionally, Solo is required to maintain LARG activity in cells, and this interaction is related to actin regulation in response to substrate stiffness. We further investigated the relationship between LARG and PRG as a function of Solo. We noted that although they did not competitively localize at Solo accumulation sites, knockdown of either PRG or LARG suppressed Solo-induced actin polymerization to the same extent as double knockdown, indicating that these signaling pathways cooperatively regulate Solo-induced actin polymerization.

## Introduction

The actin cytoskeleton is a fundamental structure that determines cell motility and morphology by forming complex structures via dynamic reorganization. Numerous actin-binding or -bundling proteins and signaling pathways are involved during the reorganization of the actin cytoskeleton (dos Remedios et al., 2003; Blanchoin et al., 2014; Pollard and Goldman, 2018). The Rho family of small GTPases, which function as switch proteins cycling between the GDP-bound inactive form and the GTP-bound active form, are key regulators of the actin cytoskeletal reorganization (Jaffe and Hall, 2005; Heasman and Ridley, 2008). Twenty members of the Rho GTPase family are encoded in the human genome. When activated, each member specifically induces a particular actin structure. Rho guanine nucleotide exchange factors (RhoGEFs) activate Rho GTPases and spatiotemporally regulate their activity in response to various external stimuli (Bos et al., 2007; Hodge and Ridley, 2016). Moreover, approximately 80 RhoGEF proteins and 70 Rho GTPase-activating proteins (RhoGAPs, inactivators of Rho GTPases) are encoded in the human genome; this diversity is thought to enable cells to induce appropriate actin structures by spatiotemporally combining the activation and deactivation of Rho GTPases (Cherfils and Zeghouf, 2013; Cook et al., 2014).

Actin cytoskeleton reorganization is vital for various physiological processes, including tissue morphogenesis and embryogenesis (Lecuit et al., 2011; Romet-Lemonne and Jegou, 2013). For instance, when subjected to tensile force from neighboring cells, cells strengthen their cell-cell adhesion sites by inducing actin polymerization and actomyosin formation. Moreover, cells induce stress fiber formation when they attach to a stiff substrate (Bachir et al., 2017). RhoGEFs regulate actin cytoskeleton reorganization in response to mechanical stress (Ohashi et al., 2024). We previously searched for RhoGEFs involved in the response of human umbilical vein endothelial cells (HUVECs) to cyclic stretch-induced reorientation. We identified Solo as a RhoGEF involved in tensile force-induced RhoA activation (Abiko et al., 2015; Fujiwara et al., 2016). Subsequently, we performed a proteomic analysis of Solo in cells using the proximity-dependent biotin identification (BioID) method (Roux et al., 2012; Branon et al., 2018), to identify Solo-interacting proteins. Notably, we identified several candidate Solo-interacting proteins, including PDZ-RhoGEF (PRG, Kunitomi et al., 2024). Upon investigating the relationship between Solo and PRG, we found that Solo predominantly regulates PRG localization. Moreover, PRG is directly activated by Solo and is required for Solo-induced actin polymerization. Furthermore, this interaction contributes to actin stress fiber formation in response to substrate stiffness. Leukemia-associated RhoGEF (LARG), a member of the same regulator of G protein signaling domain (RGS)-RhoGEF family as PRG, was also identified as a Solo-interacting protein. LARG is a RhoGEF that activates RhoA in response to a tensile force; LARG also interacts with PRG. It is involved in cancer malignancy, indicating that LARG activity could mediate the relationship between Solo and PRG. However, the function of the relationship between Solo and LARG remains unclear. Moreover, although PRG and LARG are members of the RGS-RhoGEF family, the functional differences in their interactions with Solo are unknown.

Therefore, in this study, we analyzed the effects of the interaction between Solo and LARG on actin cytoskeletal remodeling and GEF activity, and showed that Solo regulates RhoA activity by interacting with LARG through processes similar to or different from those of PRG, and is involved in the response to substrate stiffness.

## Results

### LARG localizes at Solo accumulation sites, enhances Solo-induced actin assembly, and stress fiber formation at the inner area of the basal side of cells

To investigate the interaction between Solo and LARG, EGFP-tagged Solo and mCherry-tagged LARG were expressed in Madin-Darby canine kidney (MDCK) cells, and their subcellular localization and effects on actin polymerization were observed. Solo accumulated at the basal area of the cells and induced local actin polymerization at these sites, as previously reported (Figure 1A; Kunitomi et al., 2024). LARG diffused into the cytosol and induced actin stress fiber formation in both the inner and peripheral cell regions (Figure 1, A and C). Upon coexpression with Solo, LARG translocated to Solo accumulation sites, and quantitative analysis revealed an increase in Solo-induced local actin polymerization (Figure 1, A and B). Whole-cell actin fluorescence intensity did not increase compared to that in LARG mono-expressing cells; however, the ratio of actin polymerization in the inner area to that at the cell periphery of the basal side of cells increased (Figure 1, C and D). These data suggest that Solo regulates the localization of LARG, thereby promoting actin polymerization at Solo accumulation sites and intracellular stress fiber formation.

**FIGURE 1.**
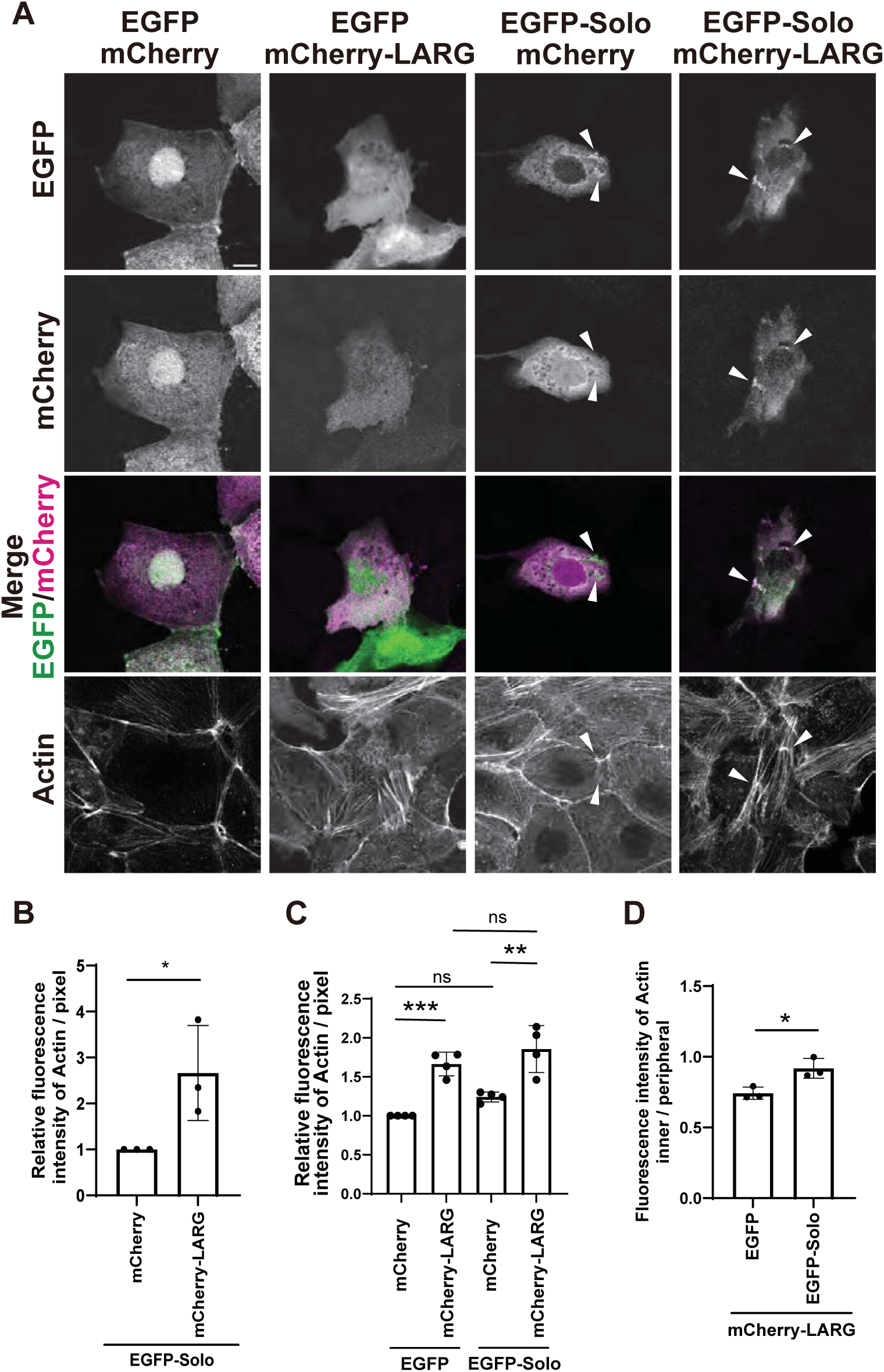
LARG colocalizes at Solo accumulated sites and induces actin polymerization at these sites and within the inner cell area. (A) Confocal microscopic images of MDCK cells transfected with the indicated plasmids. Cells were fixed and then stained with anti-mCherry antibody and Alexa 647-phalloidin. Images of each channel and merged images of EGFP (green) and mCherry (magenta) are shown. White arrowheads indicate EGFP-Solo accumulation site at the basal area of the cells. Scale bar, 10 μm. (B) Quantitative analysis of actin fluorescence intensity at Solo accumulation sites. The relative intensity is shown as the mean ± SD of three independent experiments (80-135 areas/experiment), with the intensity of cells expressing mCherry and EGFP-Solo set as 1. (C) Quantitative analysis of actin fluorescence intensity in the basal area of the cells. Actin fluorescence intensity of each cell was measured and divided by the area of the cell. The relative intensity is shown as the mean ± SD of four independent experiments (26-39 cells/experiment), with the intensity of cells expressing EGFP and mCherry set as 1. (D) Quantitative analysis of the position of actin polymerization. The ratio of actin fluorescence intensity per unit area between the inner (80% of the whole cell area) and peripheral regions (out of the inner area) of the cell was measured (see methods). Data is shown as the mean ± SD of three independent experiments (25-30 cells/experiment). ns: not significant, *p < 0.05, **p < 0.01, ***p < 0.001, (B, D two-tailed paired t-test; C one-way ANOVA followed by Tukey’ s test).

### LARG is required for Solo-induced actin polymerization

To examine whether LARG is required for Solo-induced actin assembly at Solo accumulation sites, EGFP- or EGFP-Solo-expressing MDCK cells were transfected with two different LARG-targeting siRNAs. The actin fluorescence intensity was measured in whole cells and at Solo accumulation sites. The inhibition of LARG expression was confirmed by immunoblotting (Figure 2A). Quantitative analysis of EGFP-expressing MDCK cells showed that LARG knockdown did not affect actin fluorescence intensity in whole cells (Figure 2, B and C). In contrast, in cells expressing EGFP-Solo, endogenous LARG colocalized with Solo at the basal area, accompanied by actin polymerization at these accumulation sites (Figure 2D). LARG knockdown did not inhibit the localization of Solo at the basal area of cells but decreased Solo-induced actin polymerization (Figure 2, D and E). These data suggest that LARG is required for Solo-induced actin polymerization.

**FIGURE 2.**
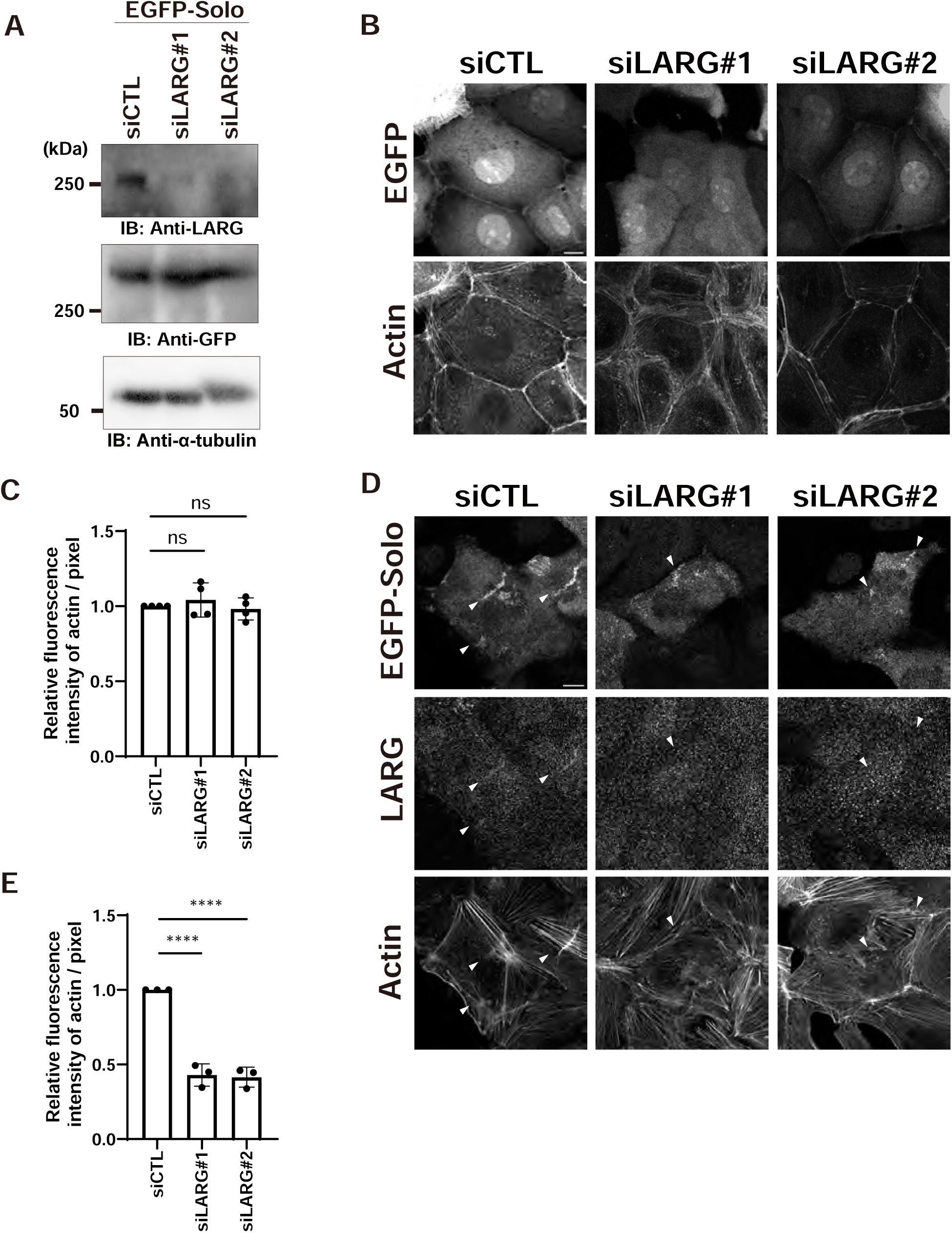
Solo-induced actin polymerization is suppressed by LARG knockdown. (A) LARG expression is suppressed by LARG-targeting siRNAs. The amount of α-tubulin in total cell lysates was used as internal control. (B) Confocal microscopic images of EGFP and F-actin in control and LARG knockdown cells. Scale bar, 10 µm. (C) Quantitative analysis of actin fluorescence intensity in the basal area of the cells. The relative intensity is shown as the mean ± SD of four independent experiments (30-36 cells/experiment), with the intensity of control siRNA transfected cells set as 1. (D) Confocal microscopic images of EGFP-Solo and F-actin in control and LARG knockdown cells. Endogenous LARG was stained with anti-LARG antibody. White arrowheads indicate Solo accumulation sites. Scale bar, 10 µm. (E) Quantitative analysis of the intensities of actin fluorescence at the Solo-accumulation sites as in Figure 1B. The relative intensity is shown as the mean ± SD of three independent experiments (118-208 areas/experiment), with the intensity of control siRNA transfected cells set as 1. ns: not significant, ****p < 0.0001 (one-way ANOVA followed by Dunnett’ s test).

### Solo is required for LARG-induced actin assembly

To investigate the role of Solo in LARG-induced actin polymerization, we examined the effects of Solo-knockdown on LARG-induced actin polymerization. The expression of Solo was reduced by siRNAs targeting Solo mRNA, whereas Solo-knockdown did not significantly affect the fluorescence intensity of F-actin in mCherry-expressing control cells (Figure 3, A and B). Additionally, Solo knockdown significantly suppressed LARG-induced actin polymerization (Figure 3C). In particular, Solo-knockdown reduced the actin fluorescence intensity in the inner cell area relative to the periphery (Figure 3D). These data suggest that Solo is required for LARG- induced actin polymerization in the inner regions of the basal area of the cell.

**FIGURE 3.**
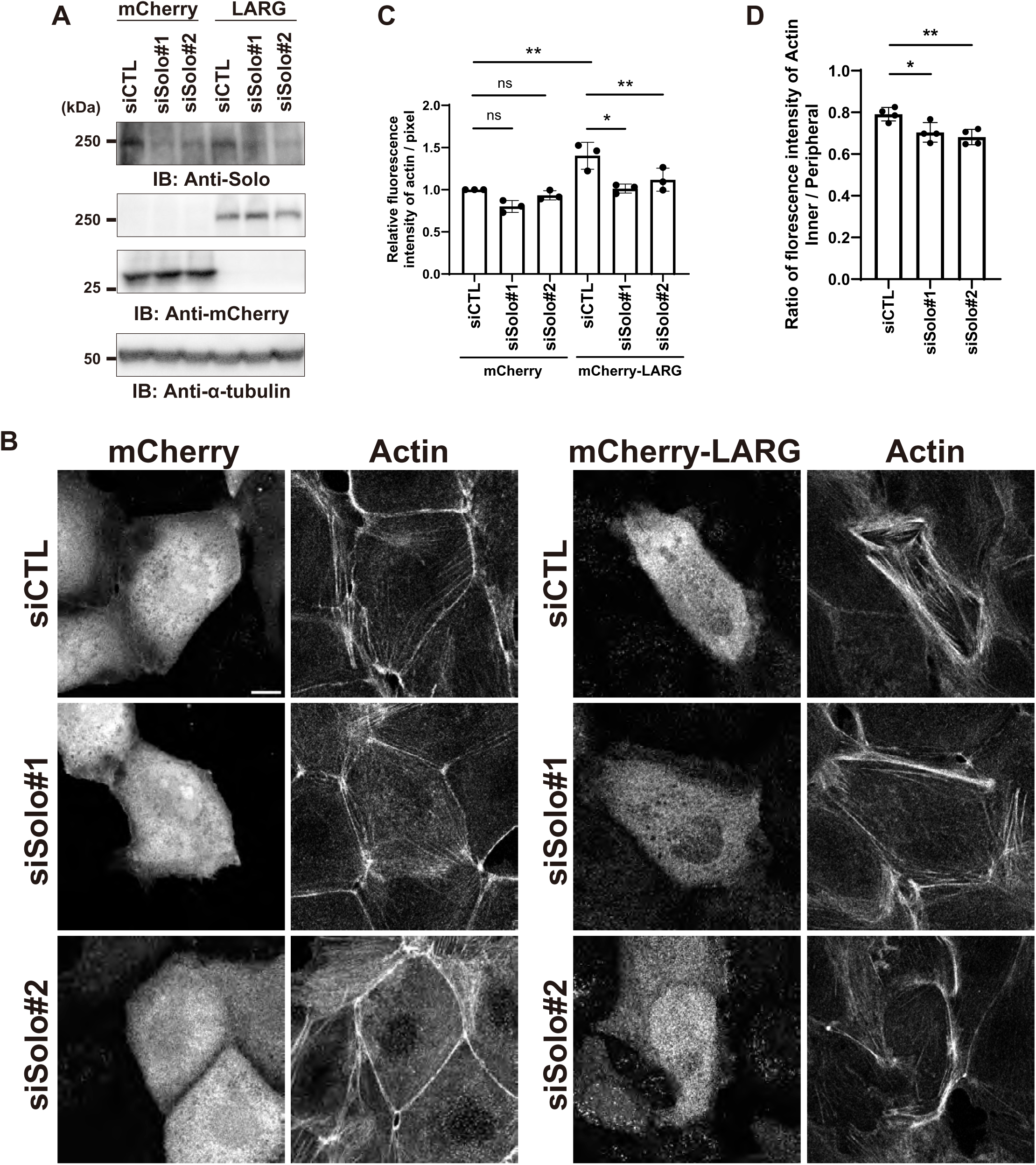
LARG-induced actin polymerization was suppressed by Solo knockdown. (A) Solo expression was inhibited by LARG-targeting siRNA. (B) Confocal microscopic images of mCherry or mCherry-LARG and F-actin in the control and Solo knockdown cells. Scale bar, 10 µm. (C) Quantitative analysis of actin fluorescence intensity in the basal area of cells as shown in Figure 1C. The relative intensity is shown as the mean ± SD of three independent experiments (28-41 cells/experiment), with the intensity of cells transfected with control siRNA and mCherry expression plasmid set as 1. (D) Quantitative analysis of subcellular position of actin polymerization as in Figure 1E. The ratio of actin fluorescence intensity of the inner to periphery is shown as the mean ± SD of four independent experiments (20-35 cells/experiment). ns: not significant, *p < 0.05, **p < 0.01 (C one-way ANOVA followed by Tukey’ s test; D one-way ANOVA followed by Dunnet’ s test).

### Mapping the binding regions of Solo and LARG and their dependence on GEF activity

LARG contains an N-terminal PDZ, RGS, and tandem Dbl homology (DH) and pleckstrin homology (PH) domains (Figure 4A; Kourlas et al., 2000). To elucidate the binding regions of Solo and LARG, we constructed LARG deletion mutants based on their respective domains. mCherry-tagged LARG or its deletion mutants and YFP-Solo WT were coexpressed in COS-7 cells and precipitated using an anti-mCherry nanobody. YFP-Solo was coprecipitated with the full-length mCherry-LARG and its deletion (N-180). The mCherry-LARG (712-C) mutants also slightly coprecipitated with YFP-Solo (Figure 4B). We also constructed a LARG GEF-inactive mutant (YA) in which Tyr-940 was substituted with Ala in the DH domain (Müller et al., 2020). An immunoprecipitation assay with YFP-Solo WT revealed that LARG YA bound to Solo WT to the same degree as LARG WT (Figure 4C). These results suggest that LARG binds to Solo via its N-terminal (N-180) region, including the PDZ domain, and 712-C fragment independent of its GEF activity. Next, to examine the LARG-binding region of Solo, COS-7 cells were transfected with YFP-tagged WT or deleted Solo, or GEF-inactive mutants, and precipitated with an anti-GFP nanobody. Solo consists of a Solo domain, a CRAL/TRIO domain, and the DH and PH domains. Deletion mutants containing each domain and a putative inactive mutant, Solo (YALELE), in which Tyr-1214 was replaced with Ala and Leu-1217/Leu-1218 with Glu in the DH domain, were used (Figure 4D; Fujiwara et al., 2016). Endogenous LARG coprecipitated with Solo WT and its deletion mutant (330–1057), and faintly with the YALELE mutant (Figure 4E). These results suggest that Solo binds to LARG through its (330–1057) region, containing the CRAL/TRIO domain, and that the GEF activity of Solo is required for stable binding.

**FIGURE 4.**
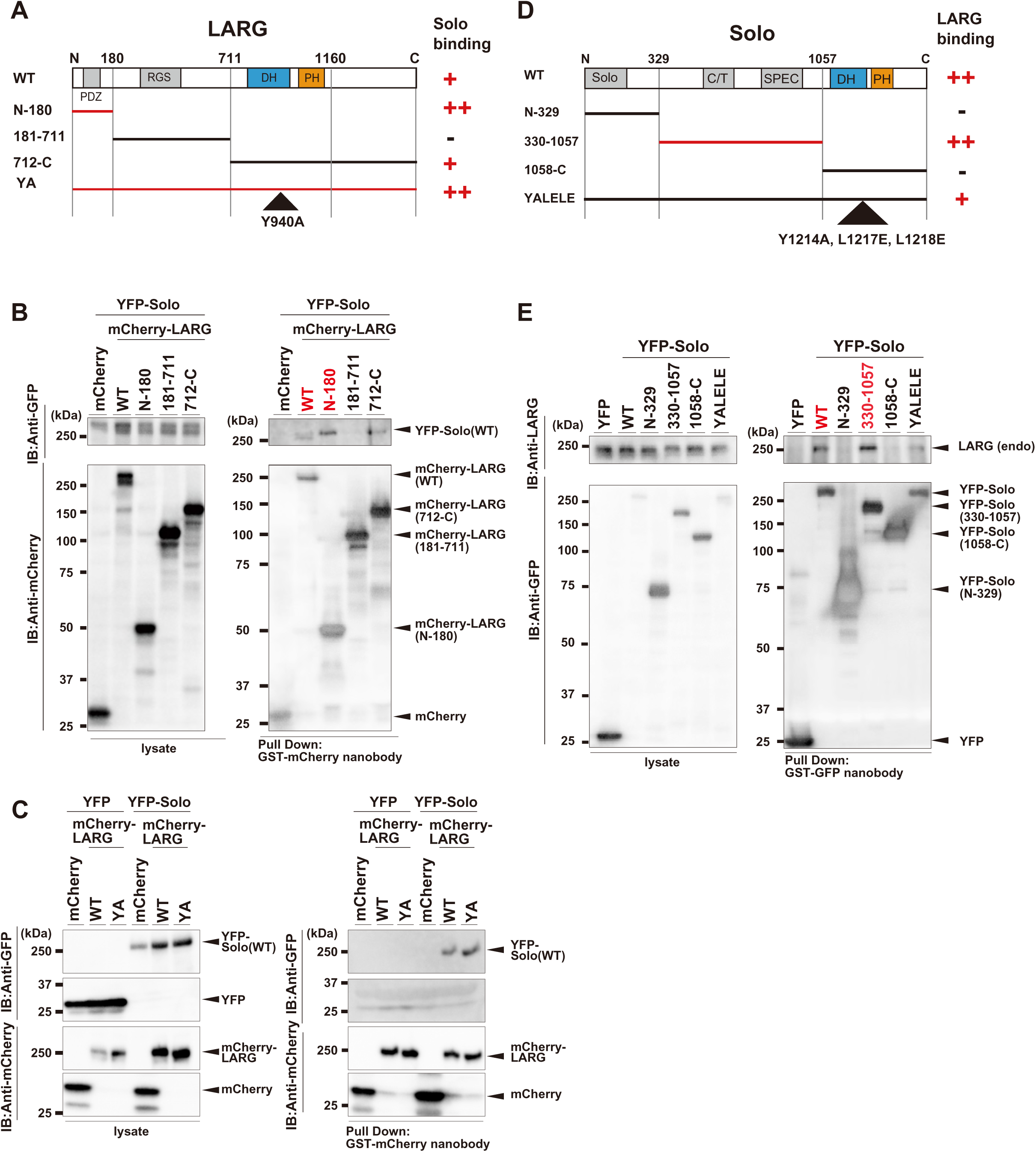
Mapping the regions required for Solo and LARG interaction in cells. (A) Schematic structures of LARG and its deletion or inactive mutants. PDZ: PDZ domain; RGS: RGS domain. (B) Analysis of the LARG regions involved in Solo binding. COS-7 cells were transfected with the indicated expression plasmids. LARG WT and its deletion mutants were precipitated from the cell lysates with anti-mCherry nanobody. The precipitated proteins were separated by SDS-PAGE and detected by immunoblotting using the indicated antibodies. (C) Coimmunoprecipitation assay using an inactive mutant of LARG (YA) with YFP-Solo (WT). mCherry, mCherry-LARG (WT), mCherry-LARG (YA) were expressed with YFP-Solo in COS-7 cells and precipitated with anti-GFP nanobody. Precipitants were analyzed by SDS-PAGE and immunoblotting as in B. (D) Schematic structures of Solo and its deletion or inactive mutants. (E) Analysis of the LARG-binding regions of Solo. Solo or its mutants were expressed in COS-7 cells and precipitated with anti-GFP nanobody. Precipitants were analyzed by SDS-PAGE and immunoblotting using the indicated antibodies.

### Solo- and LARG-induced actin polymerization requires their interaction

To investigate the effect of Solo and LARG interaction on actin polymerization in cells, we analyzed whether a dominant-negative effect was observed by overexpressing deletion mutants or GEF-inactive mutants of Solo or LARG in MDCK cells, thereby assessing their influence on LARG- or Solo-induced actin polymerization. This was done by quantifying the F-actin fluorescence intensity (Figure 5, A and B). Overexpression of the LARG deletion mutant (N-180) containing the Solo binding region significantly suppressed Solo-induced actin polymerization at its accumulation sites, but the 712-C mutant did not affect. Additionally, LARG (181–711) containing the RGS domain that binds to Gα12/13 (Booden et al., 2002), also significantly suppressed Solo-induced actin polymerization. The suppression could likely be due to the expression of LARG RGS domain exerting a dominant-negative effect on LARG GEF activity. In addition, cotransfection of MDCK cells with mCherry-LARG and Solo-deletion mutants resulted in a significant suppression of LARG-induced actin polymerization via the expression of Solo (330–1057). However, this was not observed by other Solo-deletion mutants (Figure 5, C and D). These data strongly suggest that an interaction between Solo and LARG is required for Solo-induced and partial LARG-induced actin polymerization. Subsequently, to examine whether the GEF activity of Solo or LARG was required for actin polymerization induced by LARG or Solo, EGFP-Solo or mCherry-LARG were coexpressed with the other GEF-inactive mutants, mCherry-LARG (YA) or EGFP-Solo (YALELE), respectively. The LARG (YA) mutant colocalized at Solo accumulation sites and significantly suppressed Solo-induced actin polymerization (Figure 5, A and B), whereas the Solo (YALELE) mutant did not accumulate at the basal area and did not affect LARG-induced actin polymerization (Figure 5, C and D). These results suggest that Solo induces actin polymerization using LARG as a downstream effector and that LARG executes its functions via multiple upstream signaling molecules, including Solo.

**FIGURE 5.**
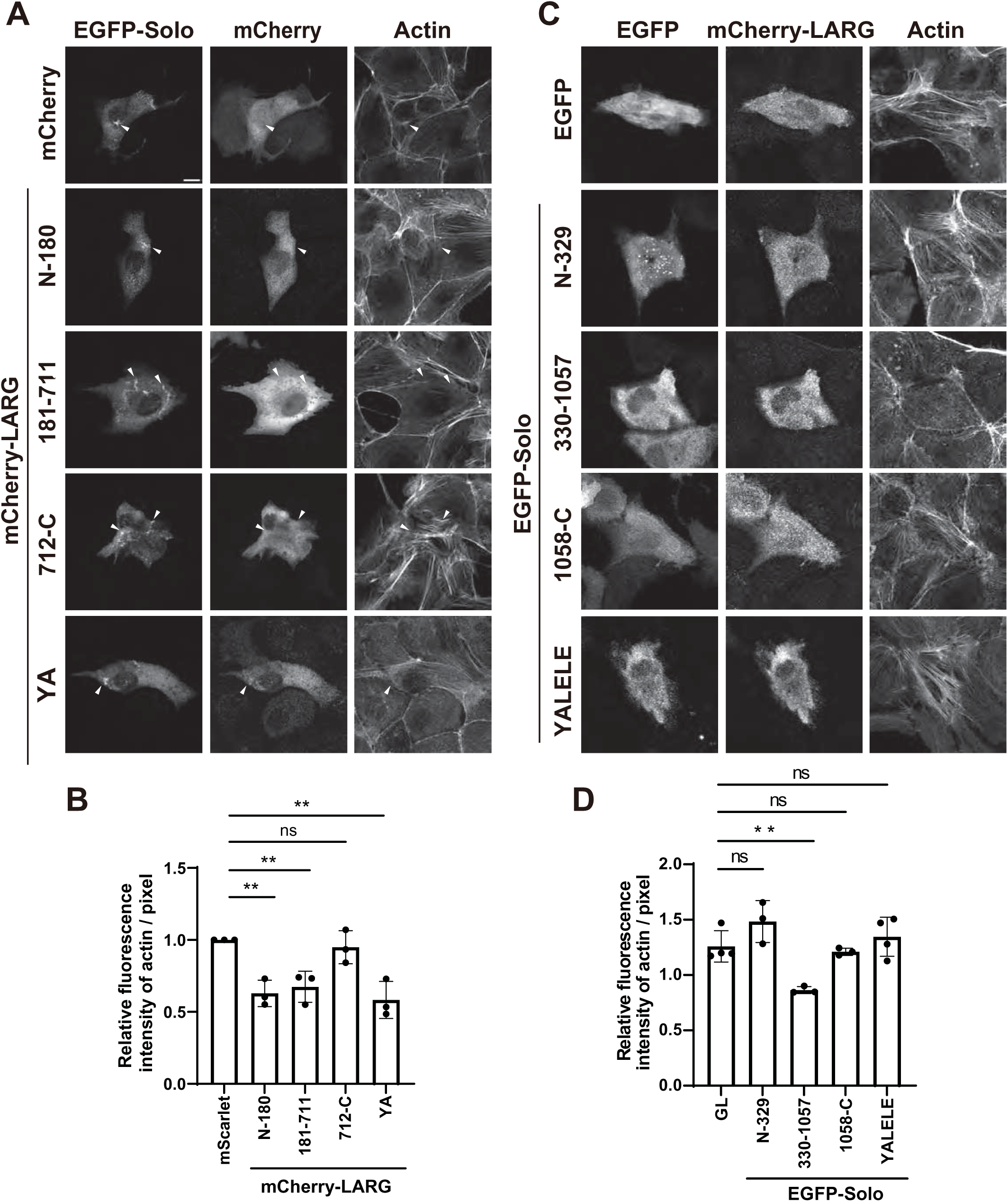
Dominant negative effects of Solo and LARG binding region overexpression on actin polymerization induced by Solo or LARG. (A) Confocal microscopic images of mCherry, mCherry-LARG, or its mutants and F-actin in MDCK cells. Cells were transfected with the indicated expression plasmids and fixed as in Figure 1A. White arrowheads indicate Solo accumulation sites. Scale bar, 10 µm. (B) Quantification of actin polymerization in Solo accumulation sites as in Figure 1B. The relative intensity is shown as the mean ± SD of three independent experiments (32-239 areas/experiment), with the intensity of mCherry and EGFP-Solo transfected cells set as 1. (C) Confocal microscopic images of EGFP, EGFP-Solo, or its mutants and F-actin in MDCK cells. Cells were transfected with the indicated expression plasmids and fixed as in Figure 1A. Scale bar, 10 µm. (D) Quantification of actin polymerization in the basal area of the cells as in Figure 1C. The relative intensity is shown as the mean ± SD of more than three independent experiments (12-29 cells/experiment), with the intensity of EGFP and mCherry-LARG transfected cells set as 1. ns: not significant, **p < 0.01 (one-way ANOVA followed by Dunnett’ s test).

### Solo- and LARG-induced RhoA activation requires their interaction

To investigate the effects of Solo and LARG interaction on their GEF activity, we performed an active RhoGEF affinity pull-down assay using a nucleotide-free RhoA G17A mutant (Gly-17 was substituted with Ala) (Garcia-Mata et al., 2006). Active LARG precipitated with glutathione-S-transferase (GST)-RhoA G17A mutant in cell lysates extracted from the control or Solo siRNA-transfected MDCK cells, revealing that endogenous active LARG was significantly reduced by Solo knockdown (Figure 6, A and B). This suggests that Solo is responsible for LARG activation in MDCK cells. Next, we examined the subcellular localization of active RhoA in YFP-Solo-expressing MDCK cells using an active RhoA probe dTomato-2xrGBD (Rhotekin G protein-binding domain) (Mahlandt et al., 2021). Although high background signal was detected due to the widespread diffusion of dTomato-2xrGBD into the cytosol, a weak fluorescent signal of the probe was detected at Solo-accumulation sites. These signals were eliminated in LARG knockdown cells (Figure 6C). These data suggest that LARG is activated by Solo and induces RhoA activation at Solo accumulation sites.

**FIGURE 6.**
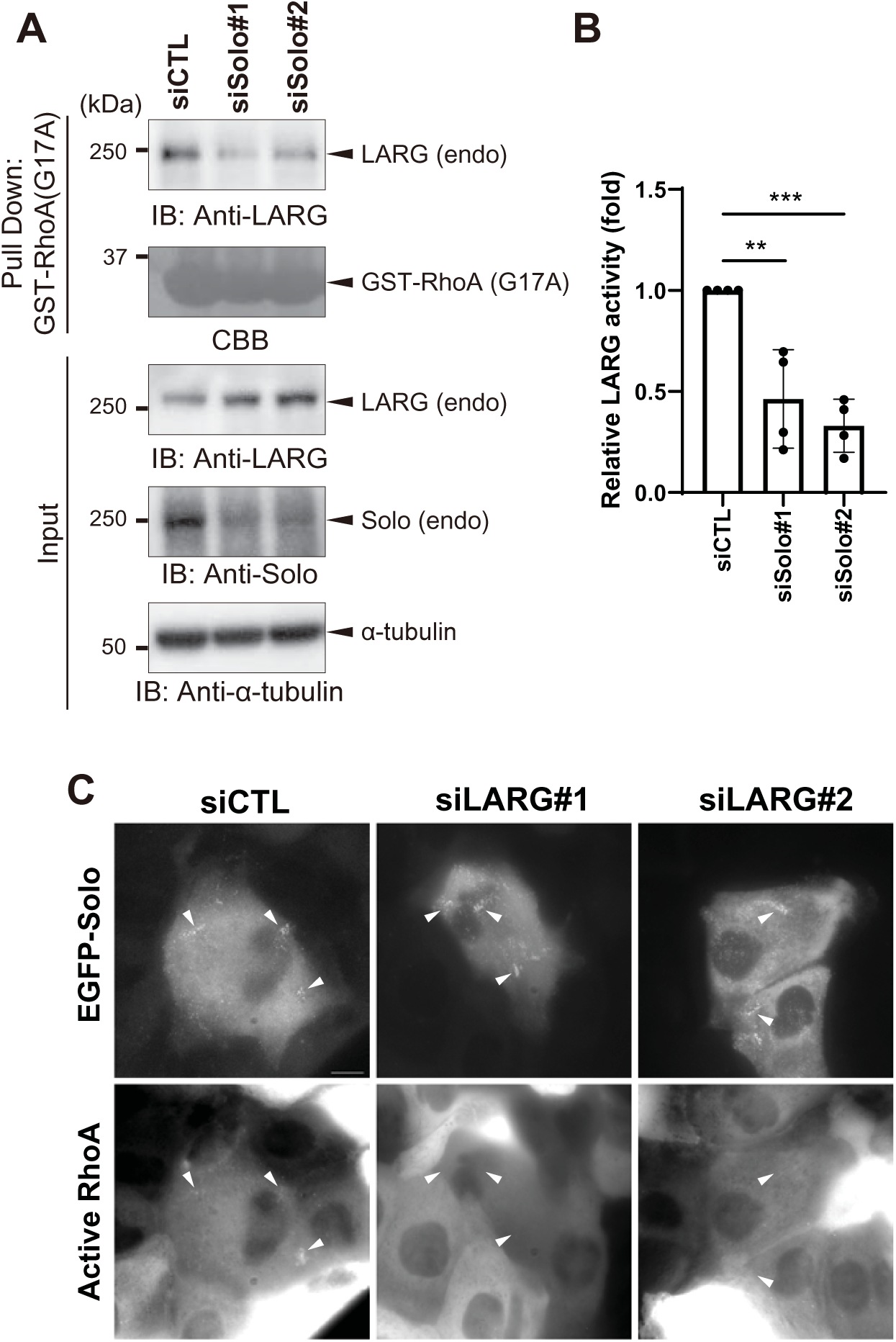
Solo is required for LARG activation in cells via RhoA activation at Solo accumulation sites. (A) LARG activity in MDCK cells transfected with control or Solo-targeting siRNAs. Cells were lysed, and the active form of LARG was precipitated with GST-RhoA (G17A). Precipitants were analyzed by immunoblotting using the indicated antibodies. (B) Quantitative analysis of GEF activity of endogenous LARG. The band signal intensity of precipitated LARG was measured and divided by the intensity of LARG in total cell lysates (input). Data is shown as the mean ± SD of four independent experiments. The activity of LARG in the cells transfected with control siRNA was set as 1. **p < 0.01, ***p < 0.001 (one-way ANOVA followed by Dunnett’ s test). (C) Fluorescence microscopic images of live MDCK cells expressing active RhoA probe, dTomato-2xrGBD and EGFP-Solo. dTomato-2xrGBD expressing cells were cotransfected with EGFP-Solo expressing plasmid and indicated siRNAs. White arrowheads indicate Solo accumulating sites. Scale bar, 10 µm.

### Solo does not directly bind to or activate LARG in vitro

To confirm whether Solo directly activates the GEF activity of LARG, we performed an in vitro exchange assay using N-methylanthraniloyl-GTP (mant-GTP), as previously reported (Kunitomi et al., 2024). The recombinant FLAG-tagged LARG protein was expressed in COS-7 cells and purified using an anti-FLAG antibody. An increase in the fluorescence of the GST-RhoA beads in the absence of GEFs was observed as the basal exchange activity of RhoA. However, GEF activity of Solo for RhoA was not observed. Meanwhile, the GEF activity of LARG on RhoA was detected (Figure 7, A and B). However, in the presence of Solo protein, the GEF activity of LARG did not increase, contrary to expectations. To examine the binding of Solo to LARG under these conditions, we performed a coimmunoprecipitation assay of Solo and LARG or PRG and found that Solo did not bind to LARG, although an interaction between Solo and PRG was detected under these conditions (Figure 7C). These data suggest the existence of another protein that mediates the interaction between Solo and LARG.

**FIGURE 7.**
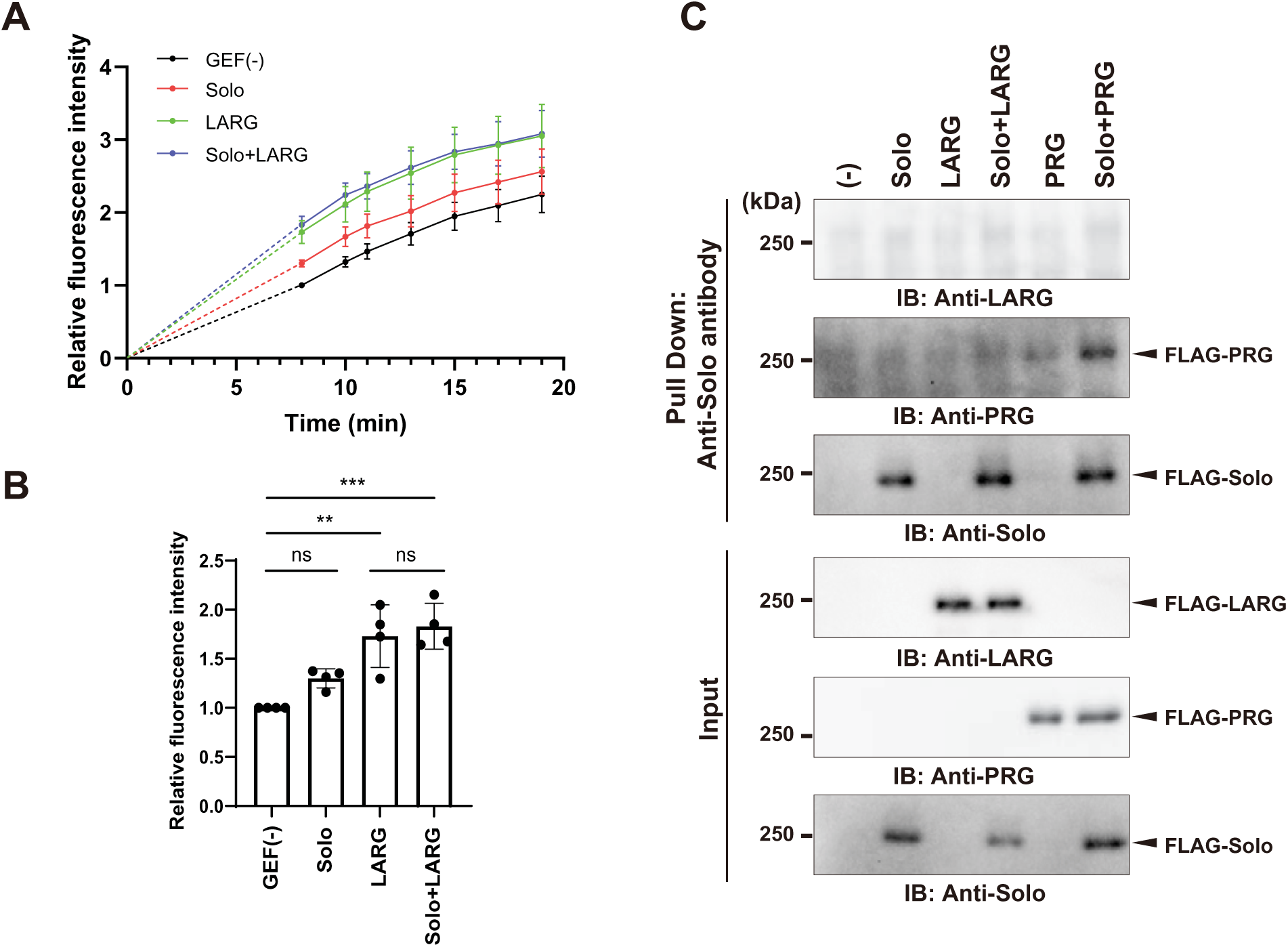
Solo and LARG do not influence each other’ s GEF activity in vitro. (A) Time course measurement of the fluorescence intensity for GDP-loaded GST-RhoA-bound beads in the GDP-GTP exchange assay for RhoA in indicated each condition. The images were acquired 8 min after the reaction started. The values are shown as the mean ± SD of four independent experiments. (B) Quantitative analysis of GEF activities. Fluorescence images of the beads were obtained, and the fluorescence intensity was measured after 8 min. Relative values are shown as the mean ± SD of four independent experiments with the value of the GEF (-) condition set as 1. ns: not significant, **p < 0.01, ***p < 0.001 (one-way ANOVA followed by Tukey’ s test). (C) Coimmunoprecipitation assay with purified GEF proteins used in A and B. The purified recombinant FLAG-tagged LARG, PRG, and Solo proteins, expressed in COS-7 cells, were mixed under the same reaction conditions as in A, and precipitated using anti-Solo antibody. Precipitants were analyzed with immunoblotting using anti-LARG, -PRG and -Solo antibodies.

### Interaction of Solo and LARG is required for substrate stiffness-dependent stress fiber formation

Solo and LARG are required for force-induced RhoA activation followed by actin polymerization and actomyosin formation (Guilluy et al., 2011; Fujiwara et al., 2016). We examined whether the interaction between Solo and LARG was involved in the mechanical stress response to substrate stiffness. MDCK cells were cultured on two substrates of different stiffness (16 and 35 kPa PA- gel), whereby the cells spread equally. The cells were then transfected with control, Solo, or LARG siRNAs, and the degree of stress fiber formation was quantified. Solo and LARG knockdown significantly suppressed stiffness-induced stress fiber formation (Figure 8, A and B). Moreover, overexpression of the binding regions of either EGFP-Solo (330-1057) or EGFP-LARG (N-180) suppressed stiffness-induced stress fiber formation (Figure 8, C and D). These findings suggest that the interaction between Solo and LARG is required for substrate stiffness-dependent actin cytoskeletal remodeling.

**FIGURE 8.**
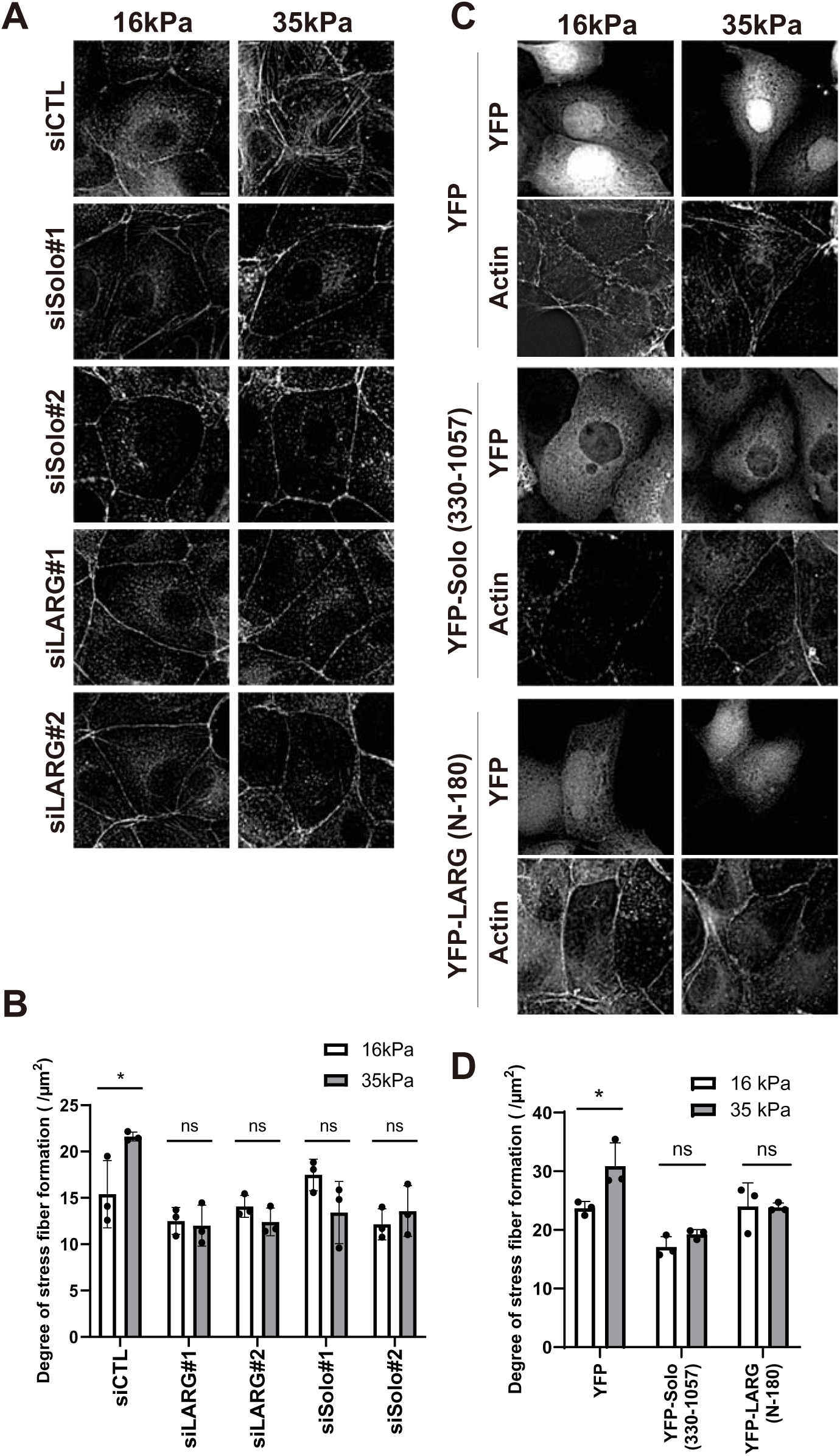
Solo and LARG interaction is involved in the stress fiber formation in MDCK cells and is dependent on substrate stiffness. (A) Deconvoluted fluorescence images of F-actin in MDCK cells transfected with the indicated siRNAs. Cells were cultured on 16 or 35 kPa PA gels cross-linked with fibronectin. Scale bar, 10 µm. (B) Quantitative analysis of stress fiber formation. Deconvoluted images were processed using the “Tubeness” plugin and “Skeletonize” function in ImageJ, followed by the acquisition of the sum intensity of Z projection images (see method). Data is shown as the mean ± SD of three independent experiments (20–43 cells/experiment). (C) Deconvoluted fluorescence images of F-actin in EGFP, EGFP-Solo (330–1057) or EGFP-LARG (N-180) expressed MDCK cells. Cells were cultured on 16 or 35 kPa PA gels, as mentioned in A. Scale bar, 10 µm. (D) Quantitative analysis of stress fiber formation as indicated in B. Data is shown as the mean ± SD of three independent experiments (11-40 cells/experiment). ns: not significant, ns *p < 0.05 (Two-way ANOVA followed by Sidak’ s test)

### PRG and LARG independently facilitate actin polymerization downstream of Solo

Our previous study showed that PRG interacts directly with Solo, localizes to Solo accumulation sites on the basal side of the cells, and facilitates actin polymerization, similar to the Solo-LARG interaction (Kunitomi et al., 2024). Subsequently, we examined whether PRG and LARG influenced each other’s Solo-dependent localization and actin polymerization. To investigate their interdependence, we examined the effect of PRG or LARG knockdown on the localization of the other. We found that their localization remained unchanged by the knockdown of the other, indicating that they localized independently at the Solo accumulation sites (Figure 9, A and B). Next, we examined the effects of the double knockdown of PRG and LARG on Solo-induced actin polymerization at its accumulation sites. The double knockdown was confirmed by immunoblotting (Figure 9C). Notably, actin polymerization at the Solo accumulation sites was not further reduced by the double knockdown compared to individual knockdowns (Figure 9, D and E). These results suggest that PRG and LARG localize at the Solo accumulation site independently, and their functions depend on each other downstream of Solo.

**FIGURE 9.**
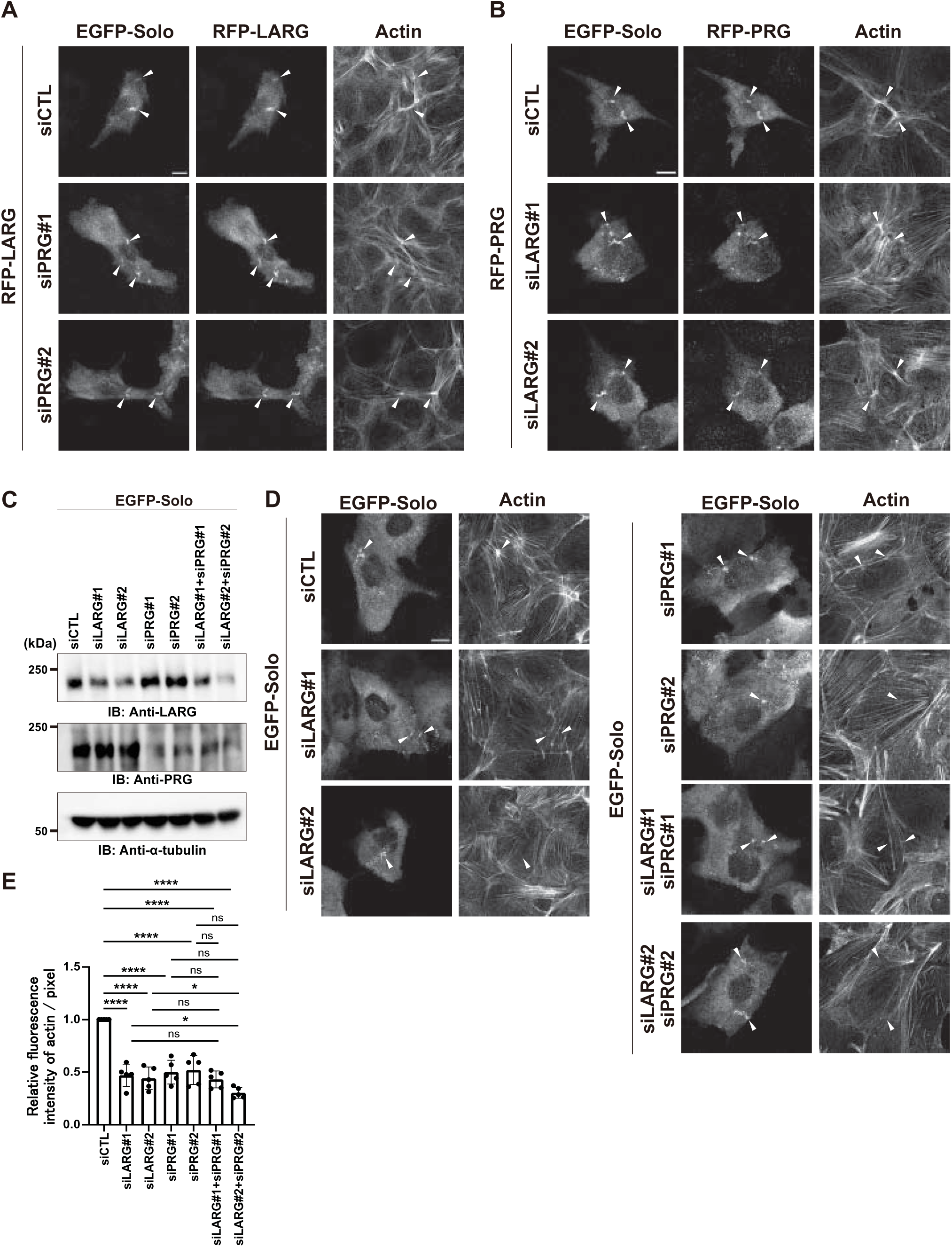
PRG and LARG localize independently at Solo accumulation sites. (A) Confocal microscopic images of F-actin, EGFP-Solo, and mCherry-LARG in MDCK cells. EGFP-Solo expressing MDCK cells were cotransfected with mCherry-LARG and control or PRG-targeting siRNAs and fixed as mentioned in Figure 1, A. Scale bar, 10 μm. (B) Confocal microscopic images of F-actin, EGFP-Solo, and mCherry-PRG in MDCK cells. EGFP-Solo expressing MDCK cells were cotransfected with mCherry-PRG and control or LARG-targeting siRNAs and fixed as in Figure 1, A. Scale bar, 10 μm. (C) Effects of siRNAs on LARG and PRG expressions. (D) Confocal microscopic images of F-actin and EGFP-Solo in MDCK cells. EGFP-Solo expressing MDCK cells were transfected with the indicated siRNA combinations and then fixed. Scale bar, 10 μm. White arrowheads in A, B, and D indicate Solo accumulation sites. (E) Quantitative analysis of actin fluorescence intensity at Solo accumulation sites. The relative intensity is shown as the mean ± SD of three independent experiments (44-85 areas/experiment), with the intensity of the control siRNA transfected cells set as 1. ns: not significant, **p < 0.01 (one-way ANOVA followed by Dunnett’ s test).

## Discussion

Our previous proteomic analysis using BioID identified LARG and PRG as candidates for Solo interaction proteins (Kunitomi et al., 2024). Comprehensive proteomic analysis of RhoGEF and LARG using BioID confirmed their interactions with Solo (O’Loughlin et al., 2018; Müller et al., 2020). Furthermore, we also revealed that PRG interacts with Solo in response to substrate stiffness to remodel the actin cytoskeleton (Kunitomi et al., 2024). However, the cellular function of the interaction between Solo and LARG remains unknown. Although Solo does not directly activate the GEF activity of LARG in vitro, unlike PRG, it is required to maintain the activity of LARG in cells. LARG is required for Solo-induced RhoA activation and actin polymerization. This interaction is involved in substrate-stiffness-dependent actin stress fiber formation. Thus, Solo appears to mediate LARG localization at its accumulation sites, where it responds to the force derived from substrate stiffness, and activates the GEF activity of LARG. In this study, we discovered another RhoGEF cascade that functions in parallel with the Solo-PRG pathway during transduction (Figure 10).

**FIGURE 10.**
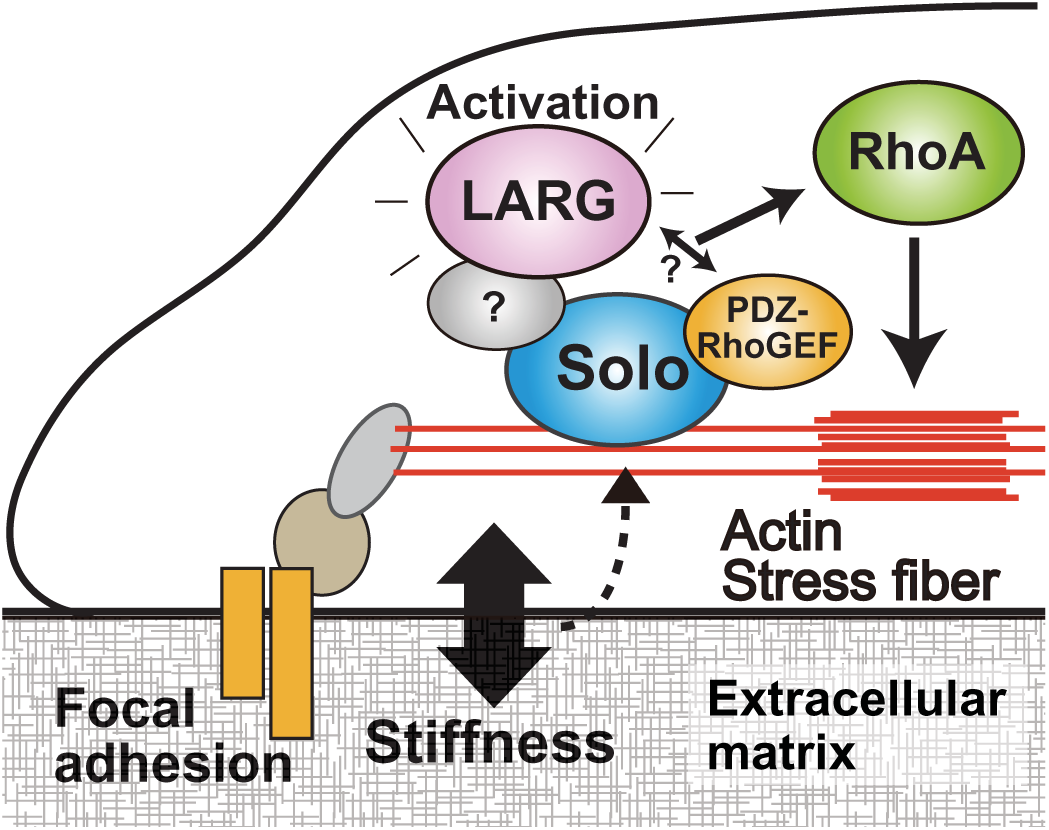
Proposed model of the cellular function Solo and LARG interaction in response to substrate stiffness and of the potential role of PRG. Solo facilitates translocation of LARG or PRG to its accumulation sites in the basal area of cells, leading to actin polymerization in a LARG- or PRG- dependent manner. Solo directly binds and activates PRG, whereas LARG is thought to be indirectly regulated by Solo through an unknown factor. Although LARG and PRG do not affect the localization of each other, they are both required for Solo-induced actin polymerization, suggesting that there may be an unknown factor that activates them and enables their localization at Solo accumulation sites.

LARG and PRG belong to the RGS-RhoGEF family, comprising the RGS domain, which is the binding domain of the α-subunit of trimeric G protein (Booden et al., 2002). PRG and LARG interact with each other and regulate cancer cell growth. Furthermore, they cooperatively regulate cell growth through RhoA activation downstream of GPCR (Chikumi et al., 2004). The present study showed that LARG and PRG both localized at Solo-accumulation sites in cells; however, their localization was found to be independent of each other. Sensing substrate stiffness is a fundamental cellular function essential for the survival of living organisms. PRG and LARG expression exhibit broad and overlapping tissue distribution in mammals. Parallel pathways are thought to ensure redundancies in substrate-stiffness sensing. Additionally, given the partially distinct roles of PRG and LARG in actin cytoskeletal remodeling (Ito et al., 2017; Castillo-Kauil et al., 2020), their involvement may vary depending on the cell type or external environment. However, our data showed that knockdown of LARG or PRG, or expression of their dominant-negative mutants, suppressed Solo-induced actin polymerization to a minimal level. Moreover, the results were not significantly different from those observed with double knockdown of LARG and PRG. Therefore, it is also speculated that LARG and PRG mutually activate each other, or that unknown factors, which can be localized by the presence of both PRG and LARG, are required to activate PRG and LARG and induce actin polymerization at the Solo accumulation sites (Figure 10). Although the LARG (712-C) mutant retains the DH-PH domain and binds to Solo, its overexpression did not enhance the actin polymerization at Solo accumulation sites (Figure 5A). This discrepancy may be explained by the fact that LARG indirectly interacts to Solo (Figure 7C) and that the 712-C mutant may have too low an affinity to colocalize with Solo (Figure 4B and Figure 5A). Further research is required to identify unknown molecules or conditions to prove the existence of molecular mechanisms.

LARG is required for RhoA activation by stimulation with lysophosphatidic acid (LPA), and we previously demonstrated that Solo is not involved in LPA-induced RhoA activation (Fujiwara et al., 2016). In contrast, LARG and Solo are required for RhoA activation when subjected to a pulling force using fibronectin-coated magnetic beads, a stimulus similar to that induced by substrate stiffness (Guilluy et al., 2011; Fujiwara et al., 2016). Therefore, it is strongly suggested that Solo and LARG interaction is involved in mechanosignals originating from integrins. Furthermore, both our study and those of others have reported that knockdown of Solo or LARG increases the velocity of collective cell migration (Medlin et al., 2010; Isozaki et al., 2020), strongly suggesting that LARG is involved in the mechanical stress response downstream of Solo.

Rho GTPases are cooperatively regulated by the interaction between RhoGEFs and RhoGAPs in cellular responses. For example, β-Pix, a GEF for Rac/Cdc42, activates Cdc42 and suppresses RhoA activity through the interaction with srGAP1, a GAP for RhoA (Kutys and Yamada, 2014). Pleckstrin homology and RhoGEF domain containing 4 B (PLEKHG4B), a Cdc42 targeting GEF, inhibits the activation of the GEF activity of LARG and PRG downstream of the G-protein-coupled receptor (Müller et al., 2020; Ninomiya et al., 2021). The interaction between Solo and PRG was the first discovery for the RhoGEF cascade that activates RhoA, followed by the interaction between Solo and LARG. Other interactions between RhoGEFs and RhoGAPs are thought to cooperatively regulate the activity of downstream RhoGTPases.

The GEF activity of Solo was not detected in the in vitro experiment, strongly suggesting that actin polymerization and RhoA activation at the Solo accumulation site are regulated by LARG. In contrast, the putative GEF inactive (YALELE) mutant of Solo did not accumulate at the basal area of the cells, suggesting that actin remodeling dependent on Solo GEF activity can trigger Solo accumulation. PLEKHG4 and PLEKHG4B, subfamilies of Solo in the Dbl family, are RhoGEFs for Cdc42 (Gupta et al., 2013; Ninomiya et al., 2021). This raises the possibility that Solo acts on a Rho GTPase other than RhoA. Alternatively, our data suggest that purified Solo does not bind to LARG in vitro, and the coprecipitation of the Solo-inactive mutant and LARG was substantially reduced. This indicates that the Solo GEF activity-induced actin structure may facilitate conformational changes in Solo to expose the LARG-binding site or may recruit an unknown factor essential for activating LARG and PRG, as mentioned above.

We also showed that the interaction between Solo and LARG is involved in actin cytoskeletal remodeling in response to substrate stiffness. Cells are thought to sense substrate stiffness by exerting tensile forces on actin filaments anchored to the cell-substrate adhesion complex, including focal adhesion. Integrin and talin, which are components of focal adhesions, function as mechanosensors that alter their conformation in response to tensile forces (Guilluy et al., 2011; Elosegui-Artola et al., 2016). Solo is localized in the vicinity of the cell-substrate adhesion complex, and integrin-mediated LARG activation requires Fyn, a member of the Src family. This suggests that the activation of Fyn at the focal adhesion site may regulate the localization and activity of Solo upstream of LARG (Guilluy et al., 2011; Lessey et al., 2012).

Solo regulates the localization and GEF activity of LARG to remodel the actin cytoskeleton; this interaction is required for cellular mechanical stress responses. However, the molecular mechanisms of Solo localization and activation by mechanical stress remain unclear, and further analysis is required.

The lack of proteins that regulate Solo-dependent actin polymerization and the characteristic localization of Solo in the proteomic analysis has stalled the functional analysis of Solo in the mechano-stress response. It may be possible to make a breakthrough by analyzing the proteome under different cell types and conditions and by examining various inhibitors. If this issue is resolved, it is expected to lead to the discovery of a novel molecular mechanism of mechanical stress responses.

## Materials & Methods

### Reagents and antibodies

F-actin was stained with Alexa Fluor 647-phalloidin (A30107, Thermo Fisher Scientific). The following primary antibodies were used: rabbit anti-LARG (sc-25638, H-70, Santa Cruz, 1:1000 for western blotting, 1:500 for immunofluorescence), mouse anti-α-tubulin (T5168, B-5-1-2, Sigma-Aldrich, 1:1000), mouse anti-GFP (632381, JL-8, Clontech, 1:1000), rabbit anti-mCherry (GTX128508, Gene Tex, 1:1000 for western blotting, 1:500 for immunofluorescence), mouse anti-FLAG (F3165, M2, Sigma-Aldrich, 1:1000 for western blotting, 1:100 for immunoprecipitation), rabbit anti-PDZ-RhoGEF (ab110059, Abcam, 1:1000 for western blotting, 1:100 for immunoprecipitation). The anti-Solo antibody was affinity-purified from rabbit antiserum raised against the C-terminal peptide (LSRQSHARALSDPTTPL) of human Solo (Abiko *et al*., 2015). The following secondary antibodies were purchased from Thermo Fisher Scientific: Alexa 568-conjugated goat anti-rabbit IgG (A11036, 1:500), horseradish peroxidase (HRP)-conjugated IgG (NA981, 1:10,000), and anti-rabbit IgG (NA934, 1:10,000).

### siRNAs

The following siRNAs using in this study were purchased from Sigma-Aldrich and used: 5’- GAGCUGAAAGAGGAACUCAAACC-3’ (dog *Solo* #1), 5’- GGGAUCAGAGACCUUUGUUUACA-3’ (dog *Solo* #2), 5’-GGAGGGAAGGAGAAUGAUA- 3’ (dog *LARG* #1), 5’-GAAGGUGAAUCGAGAUAAA-3’ (dog *LARG* #2), 5’- GAGCTCATAGAGATCCACA-3’ (dog *PDZ-RhoGEF* #1), and 5’-GGGAAAUUCUCAAGUACGU-3’ (dog *PDZ-RhoGEF* #2). The negative control siRNA was purchased from Sigma-Aldrich (MISSION siRNA Universal Negative Control).

### Plasmid construction

The cDNAs of LARG and PDZ-RhoGEF were amplified from cultured human smooth muscle cells (SMC) using PCR with PrimeSTAR HS DNA polymerase (TAKARA, Japan). Expression plasmids encoding full-length LARG or PRG, and their mutants, were amplified by PCR and inserted into mCherry-C1 (Clontech) and FLAG-C1 (Clontech). pGEX2T-GST-RhoA was generously gifted by Kozo Kaibuchi (Fujita Health University, Toyoake, Japan) and amplified using PCR to detect the G17A mutation. pGEX6P1-GFP-Nanobody and pGEX6P1-mCherry-Nanobody (LaM-2) were kindly provided by Kazuhisa Nakayama (Addgene plasmid # 61838; http://n2t.net/addgene:61838; RRID:Addgene_61838; Addgene plasmid # 162276; http://n2t.net/addgene:162276; RRID:Addgene_162276). The plasmid encoding dTomato- 2xrGBD was provided by Dorus Gadella (Addgene plasmid # 129625; http://n2t.net/addgene:129625; RRID:Addgene_129625). For retroviral plasmids, Solo-, LARG- or 2xrGBD-encoding genes were amplified by PCR and inserted into the pRetroQ vector (TAKARA). The pMD2. G plasmid was a gift from Didier Trono (Addgene plasmid # 12259; http://n2t.net/addgene:12259; RRID:Addgene_12259).

### Cell culture and transfection

MDCK, COS-7, and GP2-293 cells were obtained from Riken BRD (Japan) and cultured in Dulbecco’s modified Eagle’s medium (D-MEM) supplemented with 10% fetal bovine serum (FBS) at 37°C in a 5% CO_2_ environment. MDCK cells were transfected with expression plasmid DNAs using Lipofectamine LTX (Thermo Fisher Scientific) or siRNA using Lipofectamine RNAiMAX (Thermo Fisher Scientific). COS-7 cells were transfected with plasmid DNAs using Avalanche-Everyday Transfection Reagent (EZ Biosystems). GP2-293 cells were transfected with the retroviral plasmids using Lipofectamine 3000 (Thermo Fisher Scientific). The cell lines were checked for mycoplasma contamination every 3 months.

### Immunofluorescence staining and fluorescence imaging

MDCK cells were cultured on coverslips and transfected with a DNA plasmid or siRNA. Cells were fixed with 4% paraformaldehyde in phosphate-buffered saline (PBS) at room temperature for 20 min and permeabilized with 0.5% TritonX-100 in PBS. Then, the cells were blocked using 5% FBS in PBS, and they were exposed to primary antibodies diluted with Can Get Signal^®^ immunostain Solution A (Toyobo, Japan) overnight at 4°C. The cells were washed with PBS and exposed to secondary antibodies diluted in dilution buffer (25 mM Tris-HCl [pH 7.4], 150 mM NaCl, 0.05% Tween20, and 1 mg/mL BSA) at room temperature for 1 h, followed by washing with PBS. Images of fluorescence-stained cells were obtained using an LSM710 laser-scanning confocal microscope (Carl Zeiss) equipped with a PL Apo 63× oil objective lens (1.4 NA), a DMi8 fluorescence microscope (Leica Microsystems) equipped with a PL Apo 100× oil objective lens (1.4 NA), and an ORCA-Flash4.0 Digital CMOS camera (HAMAMATSU Photonics, Japan). Fluorescence images of cells on PA gels obtained with DMi8 were improved using a 3D non-blind deconvolution method in the LAS X software (Leica Microsystems).

### Measurement of actin polymerization

Actin polymerization was evaluated by measuring the fluorescence intensity of F-actin stained with Alexa 647-phalloidin using ImageJ software (NIH). To assess F-actin at Solo accumulation sites, confocal images of the basal plane of MDCK cells expressing EGFP-Solo were segmented using binarization using the fluorescence signal of EGFP-Solo, followed by fluorescence intensity measurements of F-actin in the segmented area. To assess F-actin in the whole cell, the cell boundaries were segmented based on the fluorescent signal of the expressed protein, followed by fluorescence intensity of F-actin within this region. To determine the ratio of F-actin intensity between the inner and peripheral cell areas, fluorescence intensity was measured inside a 20% shrunken area of cell shape and in regions between the original cell boundary and the shrunken area.

### Western blotting

Proteins were separated by sodium dodecyl sulfate-polyacrylamide gel electrophoresis (SDS-PAGE) and transferred onto polyvinylidene difluoride membranes (Immobilon, MILLIPORE). The membranes were then blocked with 5% nonfat dry milk in PBS containing 0.05% Tween-20 (PBS-T) at room temperature for 1 h. Membranes were incubated overnight at 4°C with primary antibodies diluted in Can Get Signal Solution 1 (TOYOBO) or PBS-T containing 1% skim milk. The following day, the membranes were washed thrice with PBS-T for 10 min each. Membranes were incubated with secondary antibodies diluted in PBS-T containing 1% skim milk, followed by three additional washes with PBS-T for 10 min each. Chemiluminescent signals were detected using Immobilon Western Chemiluminescent HRP Substrate (Millipore) or Immobilon ECL Ultra Western HRP Substrate (Millipore) and imaged using ChemiDoc Touch (Bio-Rad).

### Protein expression and purification of GST-GFP / mCherry nanobody

*Escherichia coli* BL21 (DE3) cells expressing either GST-GFP nanobody or GST-mCherry nanobody (LaM-2) were cultured at 37°C until they reached an optical density at 600 (OD600) of 0.5. Protein expression was induced by the addition of isopropyl β-D-1-thiogalactopyranoside (IPTG, 0.1 mM), and the culture was incubated for an additional 4 h at 30°C. Bacterial pellets obtained from 50 mL of culture were resuspended in 10 mL of binding buffer A (PBS containing 5 mM DTT and a protease inhibitor cocktail) on ice and sonicated. The lysate was treated with Triton X-100 for 30 min and centrifuged at 20,700 × *g* for 20 min at 4°C. The supernatant was incubated with 1500 µL of glutathione-Sepharose beads (50% slurry, Cytiva) at 4°C for 4 h. Finally, the GST-GFP nanobody- or GST-mCherry nanobody (LaM-2)-bound sepharose beads were washed with wash buffer A (PBS containing 5 mM DTT and 0.1% Triton X-100) and stored at 4°C.

### Coprecipitation assay

COS-7 cells (1.0×10^6^) were cultured in a 10 cm dish and transfected with the expression plasmids. The cells were then harvested with PBS and collected in a 1.5 mL tube. After centrifugation at 5400 × *g* for 3 min at 4°C, the cell pellet was lysed with lysis buffer A (25 mM Tris-HCl [pH 7.4], 140 mM NaCl, 1% TritonX-100, 2.5 mM MgCl_2_, 1 mM EGTA, 2 µg/mL pepstatin A, 10 µg/mL leupeptin, and 250 µM PMSF). The cell lysates were centrifuged at 15,000 × *g* for 10 min at 4°C. The supernatants were incubated with GST-GFP or -mCherry nanobody-bound glutathione-Sepharose beads (GFP or mCherry nanobody Sepharose) at 4°C for 4 h (Harmsen and De Haard, 2007). The beads were washed with lysis buffer A, and the precipitated proteins were eluted by boiling in Laemmli sample buffer (50 mM Tris-HCl [pH 6.8], 1.6% SDS, 8% glycerol, 20% 2-mercaptoethanol, and 0.04% BPB). The eluted proteins were then analyzed by western blotting. Coprecipitation assay was performed using a mixture of purified Solo and LARG proteins. Anti-Solo antibodies were incubated with Protein A Sepharose beads (50% slurry; Citiva) for 30 min at room temperature. The beads were then washed with binding buffer B and mixed with purified GEFs, followed by incubation at 4°C for 2 h. The beads were washed with wash buffer C (binding buffer B without BSA), and the proteins were eluted using Laemmli sample buffer. The precipitated proteins were analyzed by western blotting

### Expression and purification of recombinant GST-RhoA / RhoA(G17A) in *E*. *coli*

BL21 (DE3) cells expressing either GST-RhoA WT or G17A mutant were cultured at 37°C until they reached an OD600 and 0.5. IPTG (0.1 mM) was then added to induce protein expression, and the culture was further incubated for 16 h at 23°C. Bacterial pellets obtained after centrifugation of the cultured cells were resuspended in 20 mL of lysis buffer B (20 mM HEPES [pH, 7.5], 150 mM NaCl, 1% TritonX-100, 5 mM MgCl_2_, 1 mM DTT, 2 µg/mL pepstatin A, 10 µg/mL leupeptin, and 1 mM PMSF) on ice and sonicated. The lysate was centrifuged at 15,000 × *g* for 15 min at 4°C and the supernatant was collected. It was incubated with 500 µL of glutathione-Sepharose beads (50% slurry, GE Healthcare) at 4°C for 4 h. Finally, GST-RhoA / RhoA (G17A)-bound Sepharose beads were washed twice with lysis buffer B and wash buffer B (20 mM HEPES [pH, 7.5], 140 mM NaCl, 2.5 mM MgCl_2_, 1 mM DTT, 2 µg/mL pepstatin A, 10 µg/mL leupeptin, and 1 mM PMSF) twice, respectively, and once with wash buffer B containing glycerol and stored at 4°C.

### GST-RhoA (G17A) pulldown assay

LARG activity was detected using a pulldown assay with GST-tagged RhoA (G17A), a nucleotide-free mutant of RhoA, as previously described (Garcia-Mata et al., 2006). MDCK cells (7.5 × 10^5^) in 10 cm dishes were transfected with siRNAs. After serum starvation for 18 h, the cells were washed with PBS and lysed with lysis buffer B. Thereafter, cell lysates were centrifuged at 18800 × *g* for 5 min at 4°C, and the supernatants were incubated with GST-RhoA (G17A)-bound Sepharose beads for 45 min at 4°C. Proteins were eluted using Laemmli sample buffer and detected using western blotting. Relative LARG activity was calculated by dividing the signal intensity of active LARG by that of total LARG. The intensities of the active LARG bands were quantified using the ImageJ plug-in Band/Peak Quantification Tool (Ohgane and Yoshioka, 2019).

### Retrovirus production and infection

GP2-293 cells cultured on poly L-lysine-coated dishes were cotransfected with pMD2.G plasmid encoding VSV-G and pRetroQ vector encoding the genes of interest using Lipofectamine 3000 (Thermo Fisher Scientific). The culture medium was changed after 4 h. Supernatants containing retroviruses were collected 24 and 48 h after transfection. The collected medium was filtered using Millex-HP 0.45 µm (Millipore) and concentrated using concentrating reagent (40 mM HEPES [pH, 7.4], 400 mM NaCl, 53 mM PEG-6000) in 1/4 of the collected volume and centrifuged at 1500 × *g* for 45 min at 4°C. The pellets were resuspended in Opti-MEM and used for infection. MDCK cells cultured in D-MEM were transfected with retrovirus with 8 µg/µL polybrene.

### Active RhoA imaging

MDCK cells expressing dTomato-2xrGBD via viral gene transfer were selected using puromycin (2 µg/µL). The cells were cultured on a glass-bottom dish and cotransfected with EGFP-Solo expression plasmid DNAs and LARG-targeting or negative control siRNAs. Fluorescence signals of dTomato-2xrGBD and EGFP-Solo in live cells were observed using DMi8 microscope with a stage incubator (Tokai Hit., Co., Ltd.) set at 37°C and under a 5% CO_2_ condition.

### Expression and purification of the recombinant GEF proteins in mammalian cells

COS-7 cells (1.0×10^6^) were seeded in a 10 cm dish and transfected with expression plasmids the following day. The cells were then harvested and lysed with lysis buffer A, as described in the coimmunoprecipitation section. The anti-FLAG antibody was bound to Protein G Sepharose beads (17061801, Cytiva) by incubating for 1 h at room temperature in binding buffer B (25 mM Tris-HCl [pH 7.4], 150 mM NaCl, 0.05% Tween20, BSA 1 mg/mL) and washed with lysis buffer A. After centrifugation of the cell lysates, the supernatants were incubated with anti-FLAG antibody-bound protein G Sepharose beads (anti-FLAG antibody Sepharose) at 4°C for 2 h. The beads were washed with lysis buffer A and a FLAG-tagged precipitated proteins were eluted with the elution buffer (30 mM Tris-HCl [pH, 8.0], 50 mM NaCl, 200 µg/mL FLAG-peptide [Wako, 044-30951]) for 30 min on ice. The eluted proteins were stored at −80°C.

### GDP-GTP exchange assay

EDTA (final concentration 10 mM) and GDP (final concentration 20 µM) were added to GST-RhoA-bound Sepharose beads in the reaction buffer (20 mM Tris-HCl [pH, 7.4], 50 mM NaCl, 10 mM MgCl_2_) and incubated for 15 min at room temperature. Subsequently, MgCl_2_ (final concentration, 60 mM) was added, and the mixture was incubated for 5 min at room temperature to stop the exchange reaction. The Sepharose beads were washed and resuspended in the reaction buffer to obtain a 2.5% slurry and added to the wells of a 96-well plate (Coster, 3063). The plate then was placed on the electric stage of the DMi8 fluorescence microscope. Purified GEFs (final concentration 30 nM) and mant-GTP (final concentration 10 µM) were then added to each well. The time-lapse recording began at the specified positions with The bead images were captured every 2 min. The addition of mant-GTP and RhoGEF marked the start of the reaction. Fluorescence and DIC images were obtained using a PL Apo 20x dry objective lens (0.8 NA) and an ORCA-Flash4.0 Digital CMOS camera (HAMAMATSU Photonics). Because mant-GTP was excited at 355 nm and its emission was monitored at 488 nm, fluorescence images were obtained using a DAPI filter set. The fluorescence intensity of the beads was measured as described previously (Kunitomi et al., 2024) using ImageJ software. Regions with three or four beads in the images were selected, and the fluorescence intensity per pixel in these regions was measured.

### PA-gels fabrication

To prepare thin-layer PA gels, glutaraldehyde-treated (hydrophilized) and silicone-coated coverslips were prepared (Kunitomi et al., 2024). Acrylamide hydrogels were prepared with a bis-acrylamide to acrylamide ratio to tune stiffness (Tse and Engler, 2010). Notably, 16 or 35 kPa gels were prepared using 10:0.15% and 10:0.30% acrylamide:bis-acrylamide, respectively. Thereafter, the gels were polymerized by the addition of 0.5 µL of tetrathylethylenediamine (TEMED, Wako) and 5 µL of 10% of ammonium persulfate (APS, Wako) between the silane-coated coverslips and glutaraldehyde-treated coverslips. The gels were incubated overnight at 4°C in water. After removing the silane-coated coverslips, 1 mM sulfo-SANPAH (Pierce) was added to the gels, which were then exposed to UV light (3000 mJ /cm^2^) using a SpectroLinker XL-1500 (Spectronics Corporation). Thereafter, the gels were washed with PBS and incubated with 20 µg/mL fibronectin (Corning) in PBS. The gels were incubated overnight at 4°C and washed with PBS, and then immersed in growth medium for over 30 min. The stiffness of the gels was assessed using Young’s modulus (E) as previously described (Kunitomi et al., 2024). Briefly, a tungsten carbide sphere was placed on the hydrogels, and the depression depth was measured to fit the Hertz model. LSM710 laser-scanning confocal microscopy (Carl Zeiss) equipped with a PL Apo 10x objective lens (0.45 NA) and a DPSS 561 laser diode was used.

### Quantitative analysis of substrate stiffness-dependent stress fiber formation

MDCK cells cultured on the PA gel were fixed with 4% paraformaldehyde and stained with Alexa 647-phalloidin to visualize the actin filaments. Given that the surface of the PA-gel was bumpy, making it impossible to focus on the basal area of the whole cell in a single focal plane, the degree of stress fiber formation could not be quantified based on fluorescence intensity. To quantify the degree of stress fiber formation in response to substrate stiffness, deconvolution-processed z-slice images covering the whole cell basal area were analyzed. Deconvolved images were processed using the “Tubeness” plugin of ImageJ software to extract the line structures of actin such as stress fibers (Sato et al., 1998). The images were binarized and processed with “Skeletonize” and converted into one-pixel lines. To exclude the actin cytoskeletal structures at the cell-cell adhesion sites, the outlines of the cells at the cell-cell adhesion sites were traced. The inner area of the cell, comprising 90% of the total outline area, was designated as the measurement area. The evaluation index of the number of stress fibers was measured as the area occupied by the “one-pixel lines” per unit area.

### Statistical analysis

GraphPad Prism 10 software (GraphPad Software, La Jolla, CA) was used to generate graphics. All statistical data are expressed as mean ± standard deviation (SD) from at least three independent experiments. Comparisons between two groups were conducted using an unpaired two-tailed t-test, whereas for comparisons involving more than two groups, one-way ANOVA was applied, followed by Dunnett’s test for comparison with the control group or Tukey’s test for multiple comparisons among treatment groups. A p-value of less than 0.05 was considered statistically significant.

## ABBREVIATIONS

BioID: proximity-dependent biotin identification;
DH: Dbl homology;
GST: glutathione-S- transferase;
LARG: leukemia-associated RhoGEF;
mant-GTP: N-methylanthraniloyl-GTP;
PA: polyacrylamide;
PH: pleckstrin homology;
PLEKHG4B: pleckstrin homology and RhoGEF domain containing 4 B,
PRG: PDZ-RhoGEF;
RGS: regulator of G protein signaling domain;
RhoGEF: Rho-guanine nucleotide exchange factor

## Acknowledgments

We would like to express our gratitude to Hironori Sato (Tohoku University, Sendai, Japan) for proposing the research topic and offering insightful discussions, and Yuto Kawasaki and Koutaro Tsuru (Tohoku University, Sendai, Japan) for ther technical assistance. This work was supported by the Agency for Medical Research and Development (AMED) (grant number 18gm5810015h0003 to K.O., URL: https://www.amed.go.jp/), Advanced Graduate Program for Future Medicine and Health Care, Tohoku University, and JST SPRING (grant number JPMJSP2114 to A.K.), Additional funding was provided by JSPS KAKENHI (grant numbers JP20H03248 and JP22H05618 to K.O., and 21H02471 to K.M., URL: http://www.jsps.go.jp/j-grantsinaid/), as well as grants from the Japan Foundation for Applied Enzymology and the Uehara Memorial Foundation to K.O. A part of this study was supported by Research Equipment Sharing System, Tohoku University (041・TOF/TOF 5800 system; 090・LSM710 laser-scanning confocal microscopy). This work also used shared research equipment through the MEXT Project for promoting public utilization of advanced research infrastructure (Program for supporting the construction of core facilities; Grant Number JPMXS0440600022).

